# DrugComb - an integrative cancer drug combination data portal

**DOI:** 10.1101/560227

**Authors:** Bulat Zagidullin, Jehad Aldahdooh, Shuyu Zheng, Wenyu Wang, Yinyin Wang, Joseph Saad, Alina Malyutina, Alberto Pessia, Jing Tang

## Abstract

Drug combination therapy has the potential to enhance efficacy, reduce dose-dependent toxicity and prevent the emergence of drug resistance. However, discovery of synergistic and effective drug combinations has been a laborious and often serendipitous process. In recent years, identification of combination therapies has been accelerated due to the advances in high-throughput drug screening, but informatics approaches for systems-level data management and analysis are needed. To contribute toward this goal, we created an open-access data portal (https://drugcomb.fimm.fi) where the results of drug combination screening studies are accumulated, standardized and harmonized. Through the data portal, we provided the web server to analyze and visualize users’ own drug combination screening data. The users have an option to upload their data to DrugComb, as part of a crowdsourcing data curation effort. To initiate the data repository, we collected 437,932 drug combinations tested on a variety of cancer cell lines. We showed that linear regression approaches, when considering chemical fingerprints as predictors, have the potential to achieve high accuracy of predicting the sensitivity and synergy of drug combinations. All the data and informatics tools are freely available in DrugComb to enable a more efficient utilization of data resources for future drug combination discovery.

## INTRODUCTION

The current cancer treatment is still largely based on a “one size fits all” approach, resulting in limited efficacy due to the heterogeneity between the patients. Molecular diagnostics, histopathology and imaging techniques help stratify and monitor patients, but they provide limited support to guide treatment selection, especially for patients with recurrent cancers. NGS (Next Generation Sequencing) technologies and other omics profiling have revealed the intrinsic heterogeneity in cancer, partly explaining why patients respond differently to the same therapy (1). Even when there is an initial treatment response, cancer cells can easily develop drug resistance by the emerging activation of compensating or bypassing pathways (2). To reach effective and sustained clinical responses, many cancer patients who become resistant to standard treatments urgently need new multi-targeted drug combinations, which can effectively inhibit the cancer cells and block the emergence of drug resistance, while selectively incurring minimal effects on healthy cells (3). Although many new drugs are being developed, there is little information to guide the selection of effective combinations, as well as the identification of patients that would benefit from such combinatorial therapies. Recently, high-throughput drug combination screening techniques have been successfully applied for the functional testing of cancer cell lines or patient-derived samples, with several important hits being made (4). However, the exponentially increasing number of possible drug combinations makes a pure experimental approach quickly unfeasible, even with automated drug screening instruments (5). Therefore, data integration approaches to predict and annotate the drug combination effects at the systems level becomes a necessary route (6). To guide the patient stratification, biomarker discovery and treatment selection, a number of data harmonization, standardization and modelling challenges need to be solved before the promise of personalized drug combinations is ultimately met (7,8).

To help achieve these goals, we present DrugComb (https://drugcomb.fimm.fi/), a web-based data portal that aims to harmonize and standardize drug combination screen data for cancer cell lines. In particular, we focused on the common experimental designs where drug pairs were crossed at different doses, forming a dose-response matrix. We provided tools via a web server that allow users to visualize, analyze and annotate such dose-response data. These tools can be used for the determination of drug combination sensitivity and synergy. In addition, we provided the visualization of dose-response matrices as well as single drug response curves. Furthermore, to facilitate a crowdsourcing effort, we provided data submission tools to encourage users to share and redistribute their data in a standardized manner. Through the web server, we established a data curation pipeline to collect datasets from several major drug combination studies, covering 437,923 drug combination experiments with 7,423,800 data points across 93 human cancer cell lines. We provided the sensitivity and synergy scores for these drug combinations, and showed that these scores can be predicted by linear regression models using the structural information of the compounds. The mechanisms of action of drug combinations can be further illustrated from drug-target interaction profiles provided by major pharmacology databases including STITCH (9), PubChem (10) and ChEMBL (11). The harmonized DrugComb data can be readily linked with genomic, transcriptomic and proteomic profiles of the cancer cells, which are available in major cancer cell line databases such as COSMIC (12), CTRP (13) and MCLP (14).

DrugComb is designed to be a major source of information relevant to drug combination research, as there is currently lack of open-access services and repositories containing harmonized results of drug combinations studies. Furthermore, the analysis of drug combinations, especially in terms of their efficacy and synergy, as well as their mechanisms of action, were largely missing. With the help of data curation and analysis tools provided by DrugComb, we expect that the users may benefit from such efforts and be willing to form a community with a critical mass, so that more datasets can be collectively curated and centrally deposited. Ultimately, such a drug combination community shall lead to a consensus on the essential information that is needed to conform to the FAIR principle of research data (15). Furthermore, we expect that DrugComb will make an ideal testbed for more advanced machine learning algorithms to predict and prioritize the most effective drug combinations, which may ultimately lead to a cost-effective treatment decision support tool for the rational design of drug combinations. DrugComb prioritizes collection and dissemination of high-quality data related to drug combinations, so as to enable better understanding, validation, and prediction of synergistic drug combinations for individual cancer patients. This one-stop workflow proposed by DrugComb makes it a unique tool in cancer drug discovery research.

In this manuscript, we describe major components of DrugComb, including a web server with a variety of data analysis tools, as well as a database repository including a pipeline of how the curation and standardization of the major drug combination studies were done. Such a pipeline can be further developed into a protocol that may be adopted by a wider drug combination screen community. Furthermore, we report the initial results of the drug combination prediction as a case study, and highlight the potential of machine learning techniques to improve the efficiency of drug combination discovery. To facilitate the use of web server and the interpretation of the data analysis results, a step-by-step user guide is also provided in the Supplementary Information and will be kept up-to-date in the web site. Future aspects of DrugComb development are also discussed in Conclusions.

## DATA PORTAL COMPONENTS

The DrugComb data portal includes two major components, the web server and the database (Figure 1). The web server, mainly available at the Analysis page (https://drugcomb.fimm.fi/analysis/), consists of modules that generate the numeric and graphical results of drug combination sensitivity and synergy analyses for users’ proprietary data. The database, retrievable at the Home page, harbors the curated datasets and their analysis results that are publicly accessible. To facilitate the annotation of these drug combinations, we utilized third party APIs to access i) chemical-protein association networks in the STITCH database, ii) molecular structural information in the PubChem database and iii) ligand-based target predictions in the ChEMBL database. A registered user may also submit the proprietary data via the Contribution page (https://drugcomb.fimm.fi/contribute/), which will be evaluated by the administrator for its appropriateness to be deposited in the database. All the data visualization functionalities are built using Javascript. Computational backend employs MariaDB for the database, while R, Python and PHP routines are used for the drug combination sensitivity and synergy analyses.

**Figure 1.**
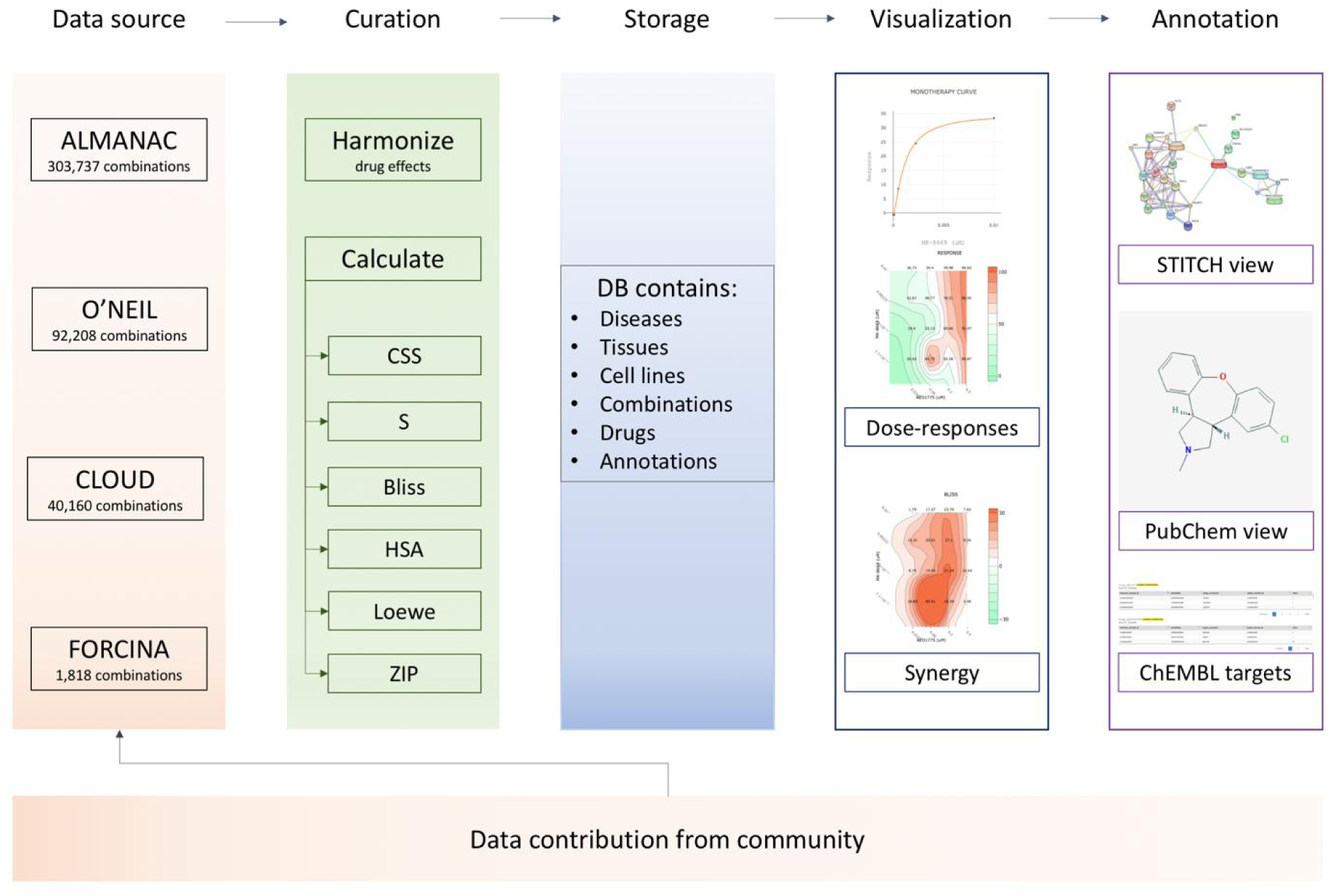
Overview of DrugComb portal and the workflow. Drug combination screen data can be uploaded by users or from the literature. Data curation includes harmonization of drug effects as percentage inhibitions compared to the DMSO negative control, and determination of drug combination sensitivity (CSS and S scores) and synergy scores (HSA, Bliss, Loewe and ZIP scores). All harmonized data and their analysis results are stored in MariaDB with regular backups, which can be visualized into dose-response curves and matrices, as well as synergy landscapes. External tools allow for network-centric representation of drug-drug interactions from STITCH database, skeletal views of drug molecules for PubChem, as well as predicted drug-target interactions from ChEMBL database.

### Computational Tools

We designed, developed and integrated a set of tools that facilitate the data processing and analysis tasks in drug combination screening research. A user needs to upload an input file that should contain information about the compounds and the cell lines, including names, concentrations and drug effects in the unit of percentage of inhibition (% inhibition) of cancer cells. Furthermore, a unique identifier, termed block id, is needed to differentiate the same drug combinations that are repeated in multiple batches, as well as serve as a unique identifier for each of the drug pairs tested. The output of the web server consists of sensitivity and synergy scores that are summarized in a table which can be further linked to more detailed graphical results. For example, the drug combination sensitivity score (CSS) is determined as the average area under the combinations’ dose-response curves with one compound fixed at the IC_50_ concentration (Unpublished material https://www.biorxiv.org/content/10.1101/512244v1, Supplementary Information). CSS summarizes the dose-responses of a drug combination screen using a metric of % inhibition, which could then be readily compared to its monotherapy drug sensitivity scores, such as DSS (16) or AAC (17). The difference between CSS and the maximal DSS of the two constitute drugs, termed as S score, is used to evaluate the benefits of a drug combination. On the other hand, to assess the degree of drug-drug interactions, also known as drug combination synergy, we provided reference models to determine the expected effect of non-interaction. Currently four commonly-used reference models were utilized, including Bliss independence (BLISS), Highest single agent (HSA), Loewe additivity (LOEWE), and Zero interaction potency (ZIP) (18–20). When two drugs are administered together their combined effect could be greater, identical or less than that predicted by their individual potencies. This is referred to as drug synergy, drug additivity or drug antagonism respectively (21). The drug combination synergy scores were then determined as the difference between the observation and expectation, with higher values being more synergistic than lower values. As these four models are based on a distinctive set of empirical or biological assumptions, which might lead to different quantification of the degree of interaction, we therefore provided all of them for users’ discretion (22). The web server also generates graphical results, including the drug combination response and synergy landscapes over the dose matrix, the monotherapy dose-response curves of its constituent drugs, and the box plots of CSS and S scores (Figure 2). The computational engine of the web server is extended from the R package synergyfinder (23), while the details on the analytical methods can be found in Supplementary Information.

**Figure 2.**
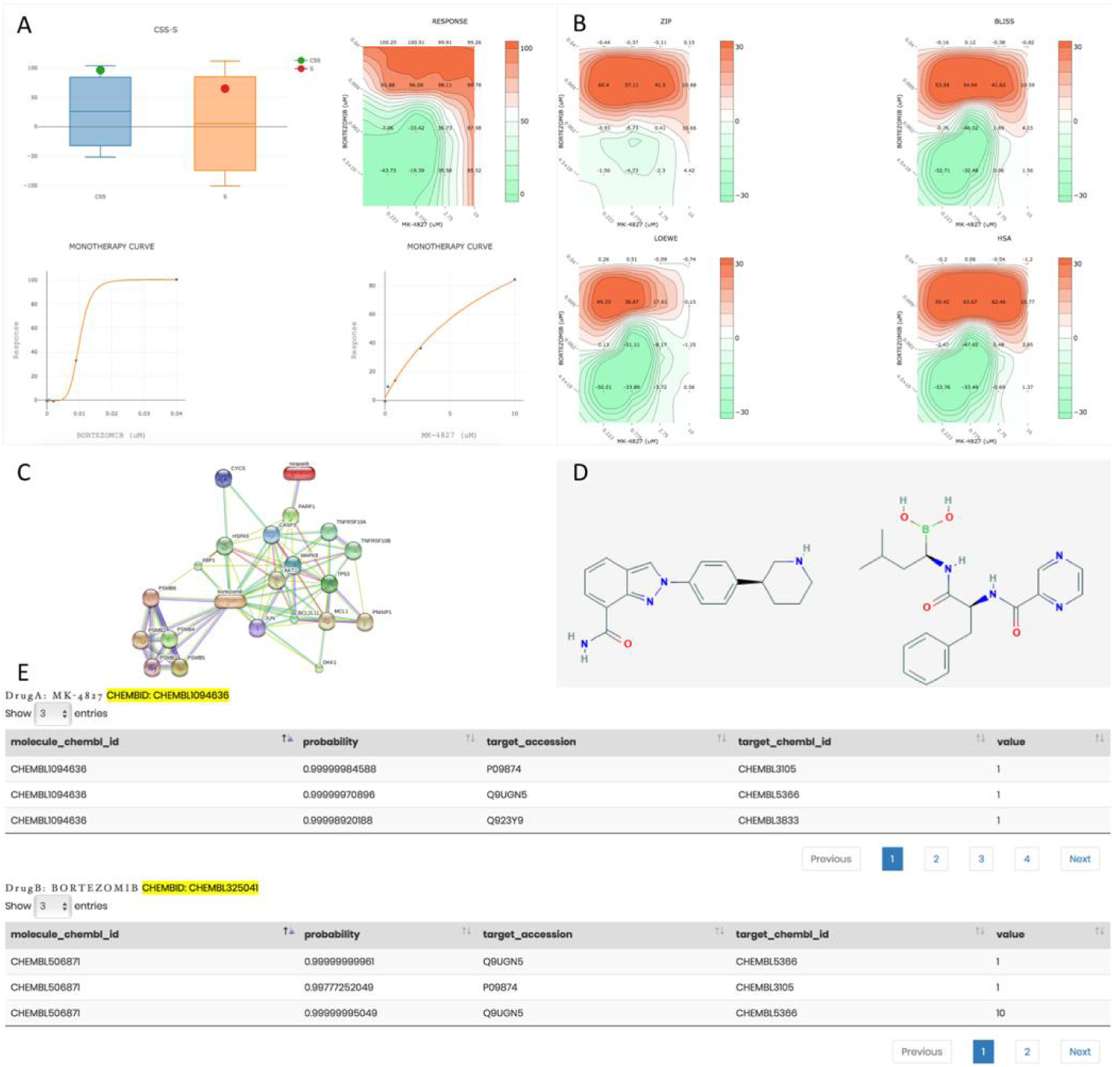
Examples of the web server analysis results, demonstrating the MK-4827 and Bortezomib combination in the MSTO cell line. (A) Single and combination dose-responses graphs, as well as the CSS-S boxplots. (B) Drug synergy landscapes determined using the HSA, Bliss, Loewe and ZIP reference models. For both panels values are colour-coded such that green corresponds to lower values and red corresponds to higher values. (C) Representation of drug-target network for the selected drug combination obtained from the STITCH database. (D) Skeletal formulae of the queried compounds from PubChem. (E) Drug-target predictions from ChEMBL.

## DATABASE CONTENT

DrugComb aims at free access to standardized drug screening results. Utilizing the computational tools that are available on the web server, we managed to collect and curate drug combination screen data involving 2276 drugs tested in 437,932 combinations for 93 cancer cell lines from 10 different tissues. The sources of the data include: i) The NCI ALMANAC dataset (24), ii) The ONEIL dataset (25), iii) The FORCINA dataset (26) and iv) The CLOUD dataset (27) (Table 1). To make the datasets comparable, we standardized the % viability values, determined as the ratio between the counts for cells treated with drugs and cells treated with DMSO as negative control, measured at the end time point. The drug effects were then represented as % inhibition values, defined as 100 - % viability. The data curation aims to determine a full dose-response matrix where the monotherapy and combination doses were matched. More specifically, in the ALMANAC dataset screenings have been performed in two different stages using two different protocols. In the first stage drugs were screened in single doses on the full NCI60 cell panel to efficiently capture compounds with anti-proliferative activity. Compounds with above threshold effects were subsequently screened in the 5-dose panel. Two different screening protocols in the second stage resulted in dose-response matrices of 6×4 and 4×4 shapes. For the ONEIL dataset the cell viability was measured as the ratio of the exponential growth rate for cells treated with a drug versus DMSO. The experiment was designed so that the monotherapy and the drug combinations were tested separately. However, the concentrations that were tested in the monotherapy screen were not identical to those in the combination screen. We thus utilized the four-parameter logistic model, available in the R drc package (28), to estimate the monotherapy responses at the concentrations tested in the combination screen. For the Forcina dataset, the % viability values were determined using the cell counts at the time of 96 hours, even though the data for other intermediate time points were also available. For the CLOUD dataset, we fitted a 4-parameter log-logistic model similar for the ONEIL dataset to estimate the % inhibition values for those drug combinations for which the single drug effects were not reported.

**Table 1.** The data statistics of the four studies curated in DrugComb.

| Study | Number of drugs | Number of drug combinations | Number of cell lines | Number of tissues | Size of the full dose-response matrix |
| --- | --- | --- | --- | --- | --- |
| **ALMANAC** | 103 | 303,737 | 60 | 10 | 4x4 or 6x4 |
| **ONEIL** | 38 | 92,208 | 39 | 6 | 5x5 |
| **FORCINA** | 1,818 | 1,818 | 1 | 1 | 2x2 |
| **CLOUD** | 283 | 40,160 | 1 | 1 | 2x2 |

For the curated drug combinations, DrugComb reported the analysis results provided by the computational tools as described earlier. Furthermore, multiple views on their annotations from other databases were also made directly available. For example, STITCH can provide a network-centric view on the drug-target interactions for a drug combination, while ChEMBL and PubChem can provide the most up-to-date information on their potential mechanisms of actions and signaling pathways, which can be further validated using experimental techniques, such as CRISPR-Cas9 or RNAi genetic screens (29,30). We provided flexible query options to navigate the repository of harmonized drug combination data and their analysis results, which may encourage users to contribute their own screening results, thus promoting a community-driven ecosystem for data sharing and redistribution. A data contribution module (https://drugcomb.fimm.fi/contribute/) is therefore provided to allow users to upload their curated datasets for which the reporting of sufficient information on the experimental procedures is mandatory.

## WEB SERVER IMPLEMENTATION

To start the DrugComb pipeline, a comma-separated values (csv) file compliant with a specific format needs to be uploaded. An example of such is provided in the Analysis page to facilitate the file generating. The server will generate the analysis outputs in two panels: Table and Graph. The Table panel is the default option which provides information about combined drugs, cell lines in which the combinations are tested, CSS as well as synergy scores determined using different reference models. The graphical results are displayed under the Graph panel, which can be activated after selecting a drug combination in the Table panel. This Graph panel contains three tabs: Sensitivity, Synergy and Annotation. The Sensitivity tab provides the results on drug combination sensitivity, including CSS-S plot, color-coded %inhibition values over the dose-response matrix, as well as monotherapy dose-response curves for the two constitute drugs. The Synergy tab contains drug combination synergy landscapes determined by the four reference models, with colour code and visualization options similar to that in the Sensitivity tab. Available only when their chemical identifiers are available, the Annotation tab contains information on the putative mechanisms of action obtained from the third-party databases including STITCH, PubChem and ChEMBL. STITCH provides drug-target interactions using evidence from experiments, databases and literature. PubChem is queried for the structural information of the drugs and ChEMBL is queried for the predicted drug targets based on the structural similarity. Predicted targets, if available, are given for each of the compounds separately Information shown in the Annotation panel should allow for further exploration of the drug-target space in a network-centric view for a selected drug combination.

DrugComb is built using PHP 7.2.11 for server-side data processing, Javascript ECMAScript 2015 for the frontend and Plotly library 1.40.0 for the generation of the interactive visualizations. Data is stored in MariaDB 10.1.37 with RMariaDB 1.0.6.9000 as the driver for interfacing with R. Software development tools including Python 3.6.7, numpy 1.14.1, pandas 0.23.4, scikit-learn 0.20.2, RDkit 2018.03.4, R version 3.5.1, synergyfinder 1.8.0 and tidyverse 1.2.1 are used in the analytical pipelines. Linux distribution CentOS-7 with the kernel 3.10.0 64-bit running on four processor cores and 64 Gb of RAM is used for hosting the web service on the in-house computational cluster. API-based access to PubChem is performed according to (https://pubchemdocs.ncbi.nlm.nih.gov/pug-rest), to STITCH using (https://www.stitchdata.com/docs/stitch-connect/api), and ChEMBL using (https://www.ebi.ac.uk/chembl/api/data/docs).

## CASE STUDIES

Here we present three case studies that have been performed on the curated data in DrugComb. The first case study involved a descriptive analysis of the dataset, where drugs and cell lines were clustered according to their mechanisms of action and tissue of origin. The second case study aimed to analyze the reproducibility of drug combination screen data. This was done via the comparison of the CSS values of replicates found across and within the study sources. The third case study employed linear regression to predict the CSS values using chemical descriptors of the drug molecules, demonstrating the potential of machine learning methods.

### Annotations of drugs and cell lines

To retrieve the mechanisms of actions of the 2,276 drugs in DrugComb, their chemical identifiers were queried from major databases including STITCH, PubCHEM, ChEMBL, DrugBank (31) and KEGG (32). These identifiers were then used for retrieving the pharmacological action information that is available in these databases. We followed the compound classification used in ChEMBL to manually determine the mechanism type, yielding the following categories with their proportions: inhibitor (28.09%), receptor (18.34%), blocker (2.98%), antagonist (2.54%), modulator (0.83%), agonist (0.79%) and activator (0.22%) (Figure 3A). In addition, 12.21% of drugs have been labeled as ‘other’ as their mechanisms of action are not common enough to be placed in new categories. Notably, the remaining 33.22% of drugs do not have well-documented mechanisms of action and hence have been labeled as ‘unknown’. To understand the mechanisms of action of these drug combinations, it becomes imperative to obtain more information on their unannotated constituent compounds. For example, MK-4541 was found in 5,772 combinations across six cancer tissues, while its pharmacology information remains unknown in those major databases. We did a literature survey and found that MK-4541 has been reported to selectively modulate androgen receptor (AR), acting as an AR agonist (33). Therefore, we expected that more compounds may be annotated similarly by searching the literature which has yet been curated. A more systematic annotation may be achieved via the DrugTargetCommons platform (https://drugtargetcommons.fimm.fi/), where the crowdsourcing efforts are utilized for extracting quantitative bioactivity values of drug-target interactions from the literature (34). For the 93 cancer cell lines, their annotations have been obtained from the Cellosaurus database (35) to determine their tissues of origin. All together 10 distinct tissues were present with lung cancer (16.13%), ovary cancer (15.05%) and skin cancer (15.05%) being the most common ones (Figure 3B). It can be seen that all the major cancer tissue types except for liver and stomach cancers are well represented in DrugComb, and thus demonstrating the general relevance of the existing data.

**Figure 3.**
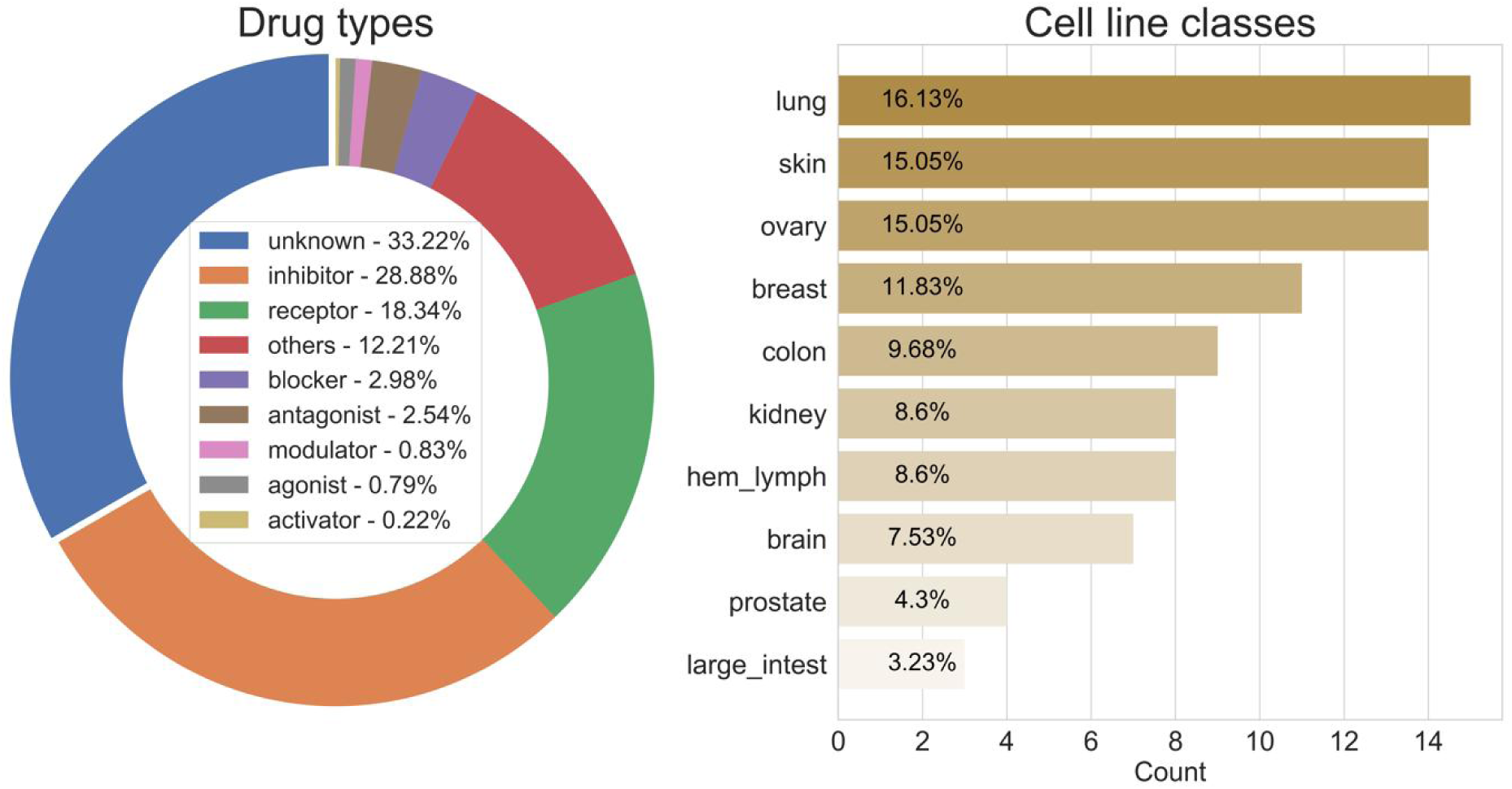
Classification of drugs and cell lines in DrugComb. Drugs were classified according to the mechanism types, following the ChEMBL implementation. Cell lines were classified according to the tissue of origin.

### Reproducibility of drug combination screens

Experimental reproducibility, in particular levels of interlaboratory concordance in the drug response phenotypes has been reported to be an issue in cancer drug screening (36). Since DrugComb aims to provide standardized results of drug combination screens, assessment of inter- and intra-study data reproducibility is of high importance. The reproducibility was evaluated using standard deviation (sd) of CSS values, which is determined for each unique drug pair and cell line combination. We chose to evaluate the CSS reproducibility as CSS indicates the average % inhibition of a drug combination and therefore makes the replicates comparable even though they were done in different concentrations. Altogether 34,936 drug-pair-cell-line combinations were replicated, while the majority of them were found either from only within the ONEIL study (n = 22,133) or from only within the ALMANAC study (n = 11,915). In contrast, the number of replicated drug combinations across the ONEIL and the ALMANAC studies is relatively few (n = 604). On the other hand, the drug combinations that were tested in the FORCINA and the CLOUD studies were not replicated, as FORCINA and CLOUD involve single cell lines of T98G and KBM-7 separately, that were not tested elsewhere. The average sd for within-study replicates is 4.25 and 12.02 for ONEIL and ALMANAC respectively, both of which are smaller than that (average sd 15.44) for their between-study replicates (p < 10_-30_, wilcoxon rank-sum test, Figure 4). The higher reproducibility of ONEIL compared to ALMANAC is expected, as the ONEIL study consisted of a standardized experiment design that involves only technical replicates while the ALMANAC study collected data from multiple labs that differed in their experimental designs, and therefore represents biological replicates in different batches (Table 1). On the other hand, for each of the n = 604 drug-pair-cell-line combinations that were replicated between ONEIL and ALMANAC, we fixed the drug-pair and picked up randomly one cell line from ONEIL and one cell line from ALMANAC, and considered the sd of the CSS values as the negative control for the between-study reproducibility. The average sd for such ‘negative control’ replicates is 17.5 which is significantly higher (p < 10_-4_, wilcoxon signed-rank paired test), suggesting a satisfactory reproducibility of the between-study replicates (Figure 4).

**Figure 4.**
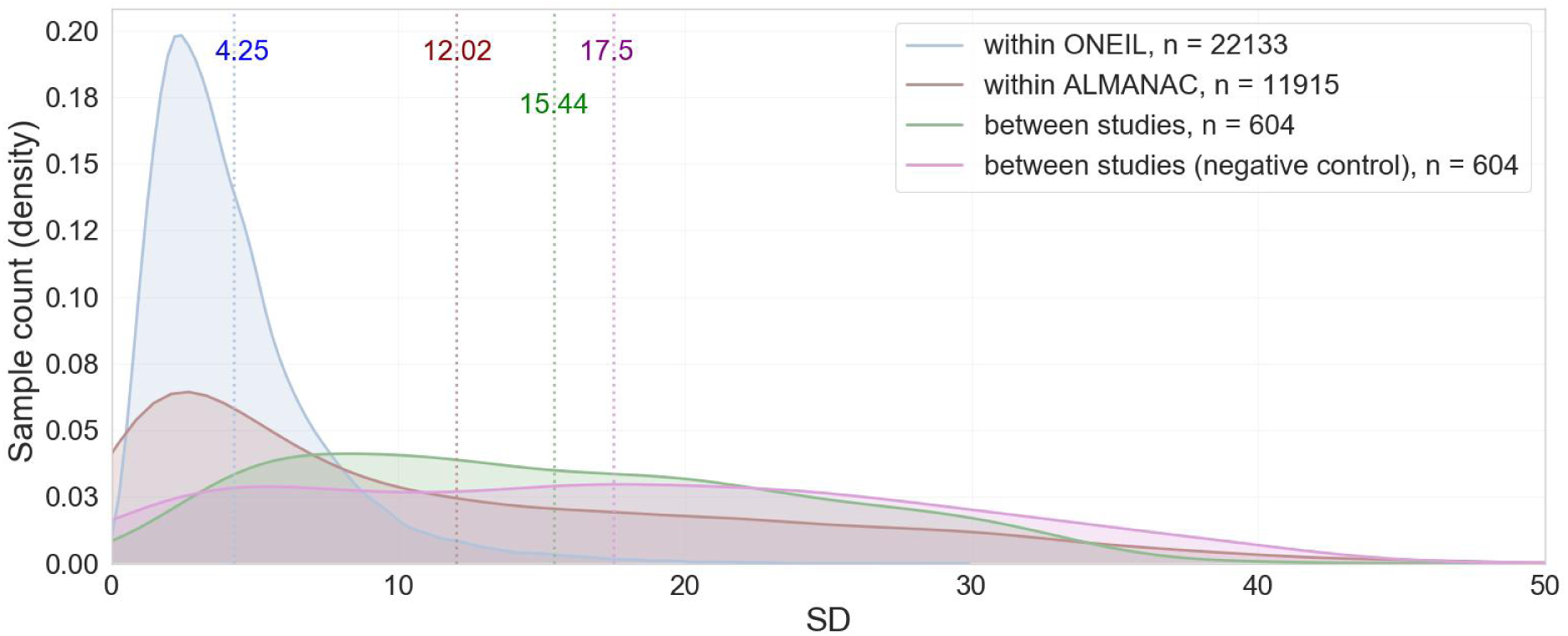
Replicability of drug combinations between and within studies represented as the distribution of the standard deviations of the Drug combination sensitivity scores (CSS). Mean values for each of the kernel density plots are delineated with a dotted line of corresponding color.

### Prediction accuracy of drug combination sensitivity

In this case study we aimed to evaluate the prediction accuracy of machine learning algorithms on the drug combination data. We considered the fingerprint information of the drug combinations as the predictors and utilized the root mean squared error (RMSE) to evaluate the prediction accuracy. To generate the fingerprint vectors for a drug combination, canonical SMILES for the constituent drugs were obtained from PubChem and then were converted to 2048 fingerprint bits using Rdkit python module (version 2018.03.4), where each bit corresponds to the presence or absence of a particular structural feature. The drug combination fingerprints were generated using the bitwise averaging of the single drug fingerprints (37). More specifically, the presence of a structural feature in both drugs yields 2 in the combination fingerprint, while presence only in one yields 1 and lack in both yields 0. These 3-bit arrays were then used as features in the machine learning algorithms. For each cell line, we fit a linear regression model on the 80% of drug combinations using a nested cross-validation and then test its prediction accuracy on the remaining 20% data. As a control, we utilized an additive model to predict CSS, which is the sum of average %inhibition from the two single drugs. The use of such an additive model was to reflect the baseline prediction assuming that the average %inhibition of a drug combination is simply the sum of their individual drug effects.

As shown in Figure 5, we found that the prediction accuracy is higher for the linear regression model than the additive model across all the tissue types, suggesting that the drug combination fingerprints carry predictive features for explaining the sensitivity. However, all the tissues exhibited multi-modality in the distribution of RMSE, suggesting a cell-line or drug-combination level heterogeneity of prediction accuracies. As a future step more advanced non-linear machine learning methods such as deep learning may be tested (38). Furthermore, molecular information of the cell lines may worth exploring for the discovery of predictive biomarkers for drug combinations.

**Figure 5.**
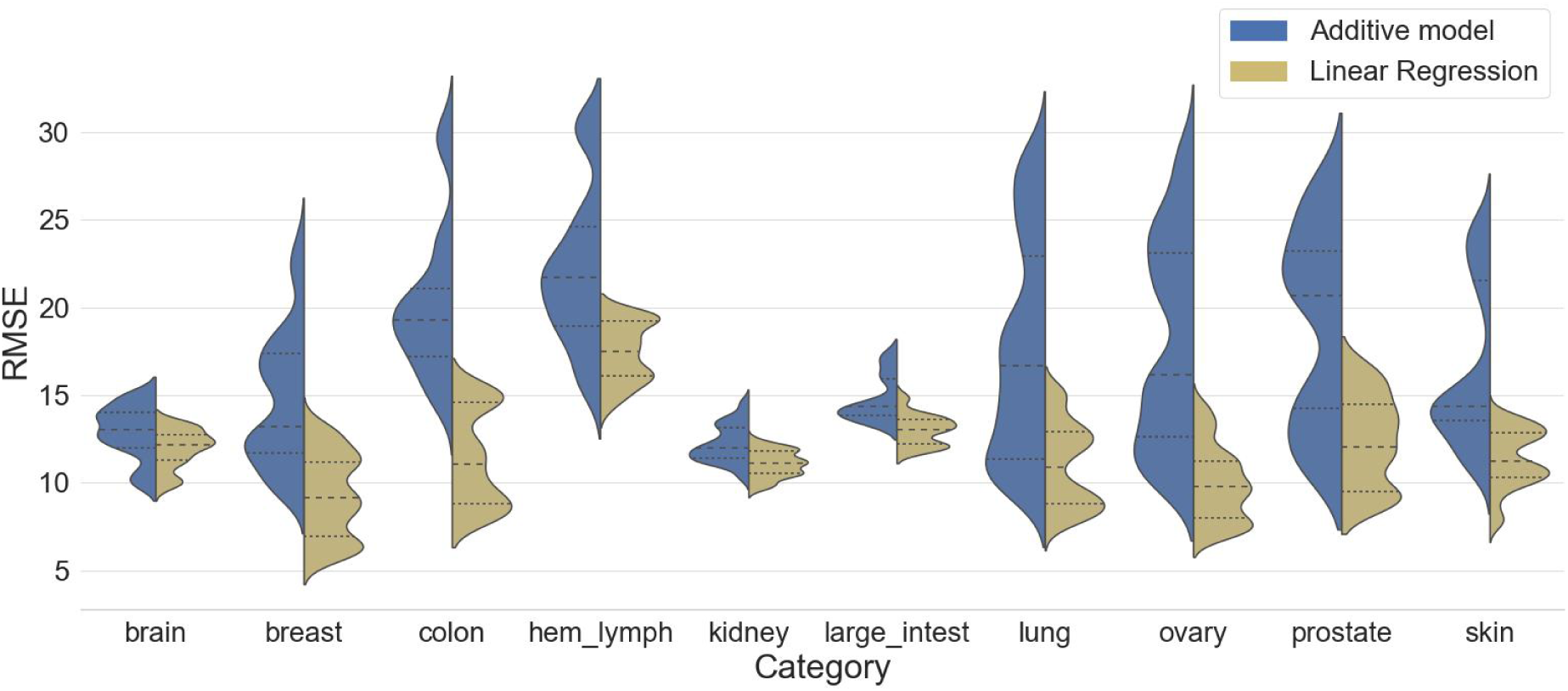
Performance of predicting CSS using linear regression as compared to the additive model. The RMSE for each cell line was grouped as density plots according to its tissue type. Dashed lines within each plot indicate interquartile range of the distribution.

## COMPARISON TO EXISTING DATA PORTALS

To the best of our knowledge, the existing data portals that cover partially drug combination screen data analysis and collection included DeepSynergy (http://shiny.bioinf.jku.at/DeepSynergy/), DrugCombdb (http://drugcombdb.denglab.org) (unpublished, https://www.biorxiv.org/content/10.1101/477547v2) and SynergyFinder (https://synergyfinder.fimm.fi/) (39). DeepSynergy provides a deep learning machine learning model that was trained on the ONEIL data and has been shown to predict new drug combinations with superior accuracy compared to conventional machine learning approaches. However, DeepSynergy did not provide the web service for the sensitivity and synergy analyses of the drug combination screen data. Furthermore, the deep learning model was trained only with the ONEIL dataset, and thus may become suboptimal when predicting a drug combination in an untested cell line. DrugCombdb is a database that harbors the concurrent screening data for 105k drug combinations. While the dataset has been collected via deep curation, it has not been analyzed with the drug combination sensitivity and synergy tools either. Therefore, both DeepSynergy and DrugCombdb provided limited web-server functionality to analyze drug combination screen data. In contrast, DrugComb provided the web-server that builds on our recent informatics approaches to assess both the sensitivity and synergy level of drug combinations, and therefore may potentially help the interpretations of the DrugCombdb data as well as contributing to the training data that is needed for DeepSyerngy and other advanced machine learning models. SynergyFinder is our recent web application for the drug combination screen data analysis. However, the focus of SynergyFinder is to analyze the degree of interactions in a drug combination screen, while the functionality of analyzing the sensitivity of drug combinations is missing. Furthermore, SynergyFinder does not provide the data curation and annotation functionality. In contrast, DrugComb provides the functionality of both a web-server and a database that have become integral components for establishing a major portal for drug combination data standardization and harmonization. Taken together, DrugComb is well positioned to provide complementary resources that can be connected with these existing tools for a more systematic and more community-driven effort for future drug combination development.

## CONCLUSIONS

How to make cancer treatment more personalized and more effective remains one of the grand challenges in the healthcare system. Drug combinations may provide enhanced efficacy to combat the cancer drug resistance and therefore may provide more sustainable treatment options for the patients. To accelerate the discovery of personalized multi-targeted drug combinations, knowledge-bases to curate, annotate and interpret the drug combination screen data are needed. The DrugComb portal provides free-access web server to analyze high-throughput drug combination screen data and thus makes it possible to develop a community-driven data repository that allows for the testing of machine learning algorithms. Future efforts include the collection of molecular profiles for cancer cell lines such that more predictive features may be extracted from the cellular genetic or epigenetic context. This may lead to the identification of biomarkers which can be used to stratify the patients for a rational selection of drug combinations. On the other hand, the curated drug combination screen data may also help define more accurate cancer cell dependency models (https://depmap.org). Furthermore, efficient statistical methods need to be developed for evaluating the significance of drug combination experimental data, which shall demonstrate that the drug combination predictions can be translated into treatment suggestions. In the long run, the DrugComb data portal is expected to provide widely applicable informatics tools to predict, test and understand drug combinations, not only for cancer cell lines but also for patient-derived samples that may lead to novel, more effective and safe treatments compared to the current cytotoxic and single-targeted therapies.

## AVAILABILITY

All the code used in generation of 3 case studies is available on github (https://github.com/netphar/first_pub)

## SUPPLEMENTARY DATA

Supplementary Data is available as a separate file and under Help section on https://drugcomb.fimm.fi

## AUTHOR CONTRIBUTIONS

JT, BZ and JA designed the study. JT and BZ wrote the manuscript. JA engineered the web server. BZ, SZ, WW, WW, YW, JS, AM, AP performed data analysis.

## ACKNOWLEDGEMENT

We thank the authors of the ALAMANC, ONEIL, FORCINA and CLOUD studies for making their drug combination data fully accessible.

